# Upregulation of eIF4E, but not other translation initiation factors, in dendritic spines during memory consolidation

**DOI:** 10.1101/2021.01.20.427437

**Authors:** Sofya Gindina, Benjamin Botsford, Kiriana Cowansage, Joseph LeDoux, Eric Klann, Charles Hoeffer, Linnaea Ostroff

## Abstract

Local translation can provide a rapid, spatially targeted supply of new proteins in distal dendrites to support synaptic changes that underlie learning. Learning and memory are especially sensitive to manipulations of translational control mechanisms, particularly those that target the initiation step, and translation initiation at synapses could be a means of maintaining synapse specificity during plasticity. Initiation predominantly occurs via recruitment of ribosomes to the 5’ mRNA cap by complexes of eukaryotic initiation factors (eIFs), and the interaction between eIF4E and eIF4G1 is a particularly important target of translational control pathways. Pharmacological inhibition of eIF4E-eIF4G1 binding impairs consolidation of memory for aversive Pavlovian conditioning as well as the accompanying increase in polyribosomes in the heads of dendritic spines in the lateral amygdala (LA). This is consistent with a role for initiation at synapses in memory formation, but whether eIFs are even present near synapses is unknown. To determine whether dendritic spines contain eIFs and whether eIF distribution is affected by learning, we combined immunolabeling with serial section transmission electron microscopy (ssTEM) volume reconstructions of LA dendrites after Pavlovian conditioning. Labeling for eIF4E, eIF4G1, and eIF2α – another key target of regulation – occurred in roughly half of dendritic spines, but learning effects were only found for eIF4E, which was upregulated in the heads of dendritic spines. Our results support the possibility of regulated translation initiation as a means of synapse-specific protein targeting during learning and are consistent with the model of eIF4E availability as a central point of control.

## Introduction

Protein synthesis in the time period immediately after learning is necessary for supporting memory-related synapse remodeling (Davis and Squire, 1984; Mayford et al., 2012; Rosenberg et al., 2014; Segal, 2016). Translation is heavily regulated, and it is now clear that learning does not simply require new proteins, but instead relies on an orchestrated balance of translational control mechanisms (Kelleher and Bear, 2008; Groppo and Richter, 2009; Darnell, 2011; Buffington et al., 2014; Santini et al., 2014). Translation occurs in neuronal processes, where it may serve as a means of spatiotemporal protein targeting at synaptic sites (Wang et al., 2010; Liu-Yesucevitz et al., 2011; Holt and Schuman, 2013; Rangaraju et al., 2017). The presence of polyribosomes and mRNA in dendritic spines suggests that local translation could be regulated at the level of individual synapses, potentially helping to maintain synapse specificity during learning-related synaptic plasticity (Steward and Levy, 1982; Tiruchinapalli et al., 2003; Kao et al., 2010; Ostroff et al., 2010; Dynes and Steward, 2012). There is evidence that local translation plays a role in learning and memory, but this comes mainly from studies of mRNA trafficking (Miller et al., 2002; Nakayama et al., 2017; Roy et al., 2020). In addition, the spatial scale of translational control in neurons is unknown (Rangaraju et al., 2017) and whether translation regulation mechanisms could mediate synapse-specific protein targeting - perhaps explaining their particular importance in learning and memory - is an open question.

The most complex regulation of translation is at the initiation step, in which ribosomes and the initiator tRNA are recruited to the target mRNA strand. Initiation is coordinated by a family of eukaryotic initiation factors (eIFs) and begins with the formation of a complex between eIF4E and eIF4G, which then binds the 5’ cap of the target mRNA and recruits a complex of the 40S ribosome and Met-tRNA bound to eIF2. The two major targets of regulation are eIF4E, whose availability to bind to eIF4G is controlled by repressor proteins, and the activity of eIF2, which is controlled by phosphorylation of its α subunit (Groppo and Richter, 2009; Sonenberg and Hinnebusch, 2009; Borden and Volpon, 2020). Dysfunction of eIF4E or eIF2α leads to impairment of various forms of learning and learning-related synaptic plasticity (Gkogkas et al., 2010; Buffington et al., 2014; Santini et al., 2014) as well as intellectual functions in humans (Kelleher and Bear, 2008; Darnell, 2011; Kapur et al., 2017). A variety of mechanisms are known to transport mRNA into dendrites and maintain it in a dormant state (Hutten et al., 2014; Buxbaum et al., 2015), and a single synapse could be the locus of translational control if initiation occurs on dormant mRNA in dendritic spines. Consistent with this possibility, we have found by serial section transmission electron microscopy (ssTEM) reconstruction that polyribosomes are upregulated in dendritic spines of the rat lateral amygdala (LA) during consolidation of aversive Pavlovian conditioning (Ostroff et al., 2010), and that the drug 4EGI-1, which inhibits initiation by interfering with the eIF4E-eIF4G interaction, impairs both formation of the memory and the associated polyribosome upregulation (Hoeffer et al., 2011; Ostroff et al., 2017).

Initiation at individual synapses is not the only explanation for this, however. Ribosomes can be stalled on mRNAs after initiation, and most dendritic polyribosomes in cultured neurons in fact appear to be stalled (Richter and Coller, 2015; Langille et al., 2019). Meanwhile, there is also evidence that actively translating mRNAs are mobile in cultured dendrites (Wu et al., 2016). Polyribosomes, either stalled or active, could be transported into spines by mRNA trafficking mechanisms, making mRNA transport, not initiation, the key regulator of synapse-specific protein targeting. To determine whether individual synapses possess initiation machinery, we combined ssTEM with immunohistochemistry for eIFs in LA dendrites following Pavlovian conditioning. We found that eIFs are indeed present in dendritic spines, and that their distribution after learning is consistent with cap-dependent initiation in spines.

## Materials and Methods

### Subjects and behavior

Adult male Sprague-Dawley rats (Hilltop) weighing approximately 300g were housed singly on a 12-hour light cycle with food and water ad libitum. Experiments were conducted during the animals’ light cycle. Behavior experiments were conducted as previously described (Ostroff et al. 2010). Briefly, animals were habituated on two consecutive days to a lit training chamber equipped with a speaker and a grid floor. On the third day, one group of animals were presented with five auditory tones (30 s, 5 kHz, 80 dB) co-terminating in a mild footshock (0.7 mA, 1 s), while a control group was placed in the chamber but not presented with tones or shocks. All procedures were approved by the Institutional Animal Care and Use Committee of New York University.

### Antibodies

The following antibodies and dilutions were used: Bethyl Labs rabbit polyclonal anti-eIF4E (A301-154A; lot# A301-154A-1; RRID:AB_2097708) at 1:500, Abnova mouse polyclonal anti-eIF4G1 (H00001981-A01; lot# 08213-2A9; RRID:AB_462490) at 1:500; Cell Signaling mouse monoclonal anti-eIF2α (L57A5; lot# 3; RRID:AB_836874) at 1:500; Vector Laboratories biotinylated polyclonal goat anti-rabbit (BA-1000; RRID:AB_2313606) or goat anti-mouse (BA-9200; RRID:AB_2336171) secondary antibodies at 1:200. Preadsorption controls were run for the two polyclonal primary antibodies by incubating them for 1 hour in a 10:1 (by weight) excess of immunizing peptide before use. Preadsorption eliminated 98% of labeled structures, as described previously (Ostroff et al., 2019).

### Immunolabeling and electron microscopy

For eIF4E and eIF4G1 labeling, rats were deeply anesthetized with chloral hydrate and perfused transcardially with ~50 ml of heparinized saline (154mM NaCl/2mM CaCl_2_/4mM MgCl_2_ in 0.01M PIPES buffer at pH 7.4) followed by 500ml of perfusion fixative (0.5% gluteraldehyde/4% paraformaldehyde/2mM CaCl_2_/4mM MgCl_2_ in 0.1M PIPES buffer, pH7.4). For eIF2α labeling, the heparinized saline was 0.01M phosphate buffer with 154 mM NaCl, and the perfusion fixative was 0.25% glutaraldehyde/4% paraformaldehyde in 0.1 M phosphate buffer, pH 7.4. All fixation and labeling procedures were carried out at room temperature. Brains were rinsed in buffered saline (0.01 M fixation buffer with 154 mM NaCl) and sectioned coronally at 40um. Sections from the left hemisphere containing the caudal LA were reacted for 30 minutes with 1% sodium borohydride then blocked in 1% BSA in buffered saline and incubated overnight in primary antibody in 1% BSA. Labeling for eIF4E and eIF4G1 was performed on alternating sections from the same rats. Sections were incubated for 30min in secondary antibody followed by 30 min in Vector Laboratories ABC-peroxidase (PK-6100; RRID:AB_2336819), then reacted with 1mM 3,3’-diaminobenzidine in 0.003% H2O2 for 5 min. The LA was dissected out of each section and processed for serial EM as previously described (Ostroff et al., 2010). Briefly, sections were postfixed in reduced osmium (1.5% potassium ferrocyanide/1% osmium tetroxide) followed by 1% osmium tetroxide, dehydrated in ethanol containing 1.5% uranyl acetate, and embedded in epon resin (LX-112, Ladd Industries). Serial 45 nm sections were cut on an ultramicrotome (Leica) in the ventral-to-dorsal direction, perpendicular to the plane of vibratome sectioning. Sections were picked up on pioloform-coated slot grids and imaged at 100kV on a JEOL 1200EX II electron microscope with an AMT digital camera. Grids were not counterstained to avoid the possibility of staining artifacts obscuring immunolabel.

### Reconstruction and analysis

Image alignment, reconstruction, and measurements were done using Reconstruct software (Fiala, 2005; RRID:SCR_002716) with experimenters blind to experimental condition. Series consisted of an average of 99 ± 4 sections. All dendrites on the central section of each series were examined and segments of spiny dendrites with a diameter less than 1μm (average 0.68 ± 0.02 μm) and length between 2 and 7 μm (average 3.87 ± 0.12) were reconstructed. There were no significant differences between antibodies or training groups in either measure. Frequencies were calculated using inclusion lengths measured from the first complete protrusion at the ventral end of the series to the final protrusion at the dorsal end. Synapse morphology was designated as asymmetric (putatively excitatory) or symmetric (inhibitory or modulatory) according to established morphological criteria (Peters et al., 1991). Approximately 11% of dendritic protrusion origins gave rise to more than one protrusion. These branched protrusions did not differ in any measures from protrusions arising singly, thus branches were treated as single protrusions for analysis. Approximately 2% of dendritic protrusions carried more than one asymmetric synapse, 4% carried both a symmetric and an asymmetric synapse, and 4% lacked a synapse. Spines were defined as protrusions carrying at least one synapse, and protrusions lacking synapses were excluded from spine analyses.

### Statistics

Analyses were conducted with STATISTICA software (Tibco Inc.). Means were compared using hierarchical ANOVAs with subjects nested into a training group to test for significant interactions. The breakdown of dendrite and spine numbers by rat, group, and antibody are given in Table 1. Bar graphs show means and standard errors, and exact p, F, and η^2^ values are given in Table 2.

**Table 1.**
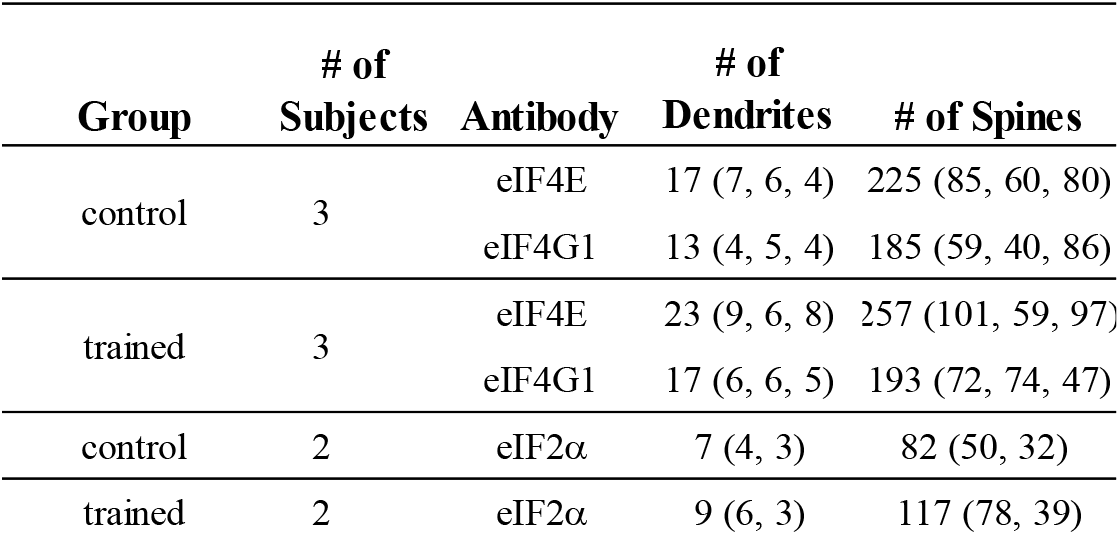
Composition of dataset. Numbers of dendrites and spines analyzed by subject, group, and antibody with individual subjects given in parentheses.

**Table 2.**
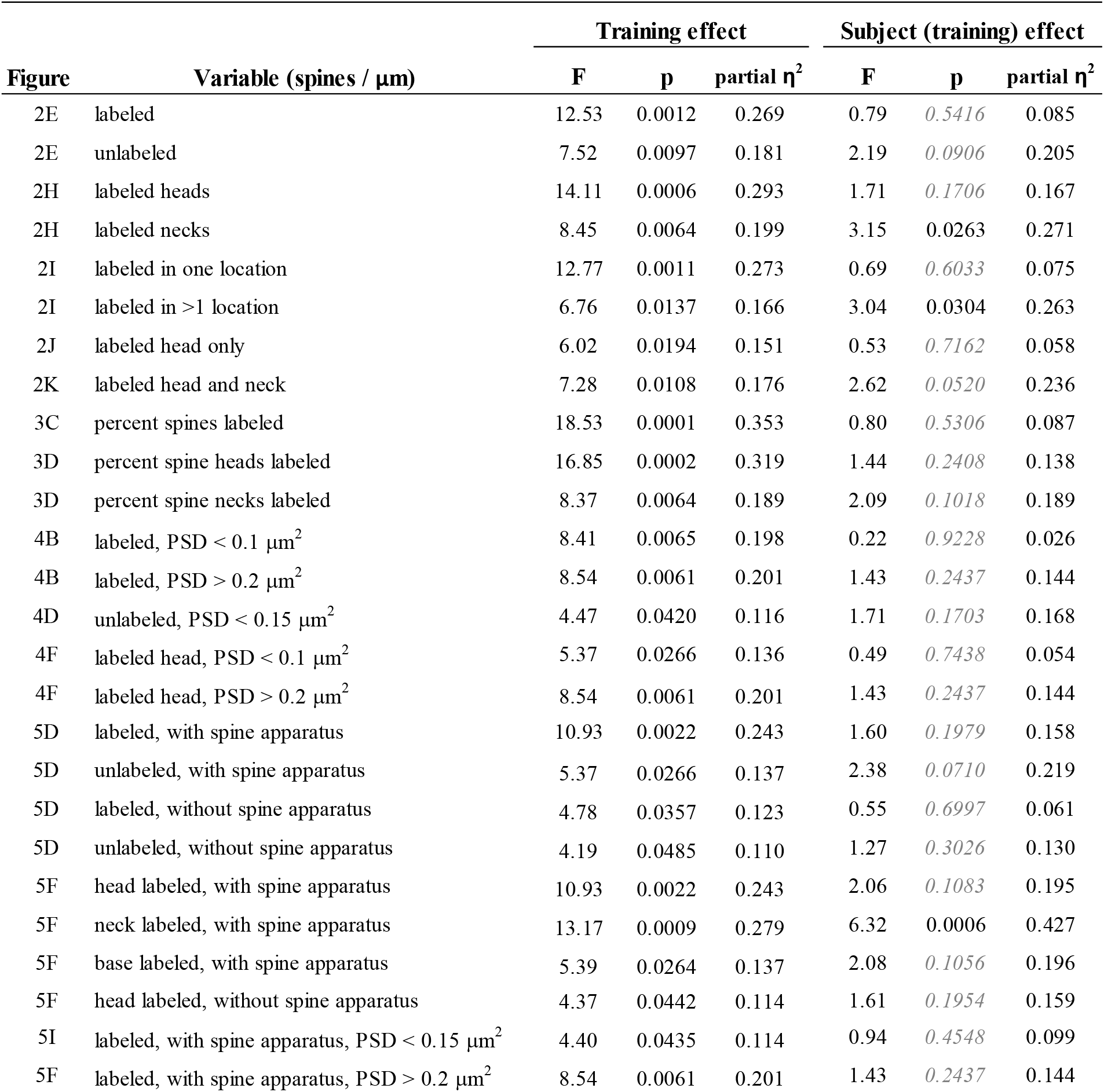
Effects of training on eIF4E labeling. Results of ANOVAs for effects shown in Figures 2-5. p values > 0.05 shown in italics.

| Figure | Variable (spines / $\mu\text{m}$) | Training effect | | | Subject (training) effect | | |
| --- | --- | --- | --- | --- | --- | --- | --- |
| | | F | p | partial $\eta^2$ | F | p | partial $\eta^2$ |
| 2E | labeled | 12.53 | 0.0012 | 0.269 | 0.79 | *0.5416* | 0.085 |
| 2E | unlabeled | 7.52 | 0.0097 | 0.181 | 2.19 | *0.0906* | 0.205 |
| 2H | labeled heads | 14.11 | 0.0006 | 0.293 | 1.71 | *0.1706* | 0.167 |
| 2H | labeled necks | 8.45 | 0.0064 | 0.199 | 3.15 | 0.0263 | 0.271 |
| 2I | labeled in one location | 12.77 | 0.0011 | 0.273 | 0.69 | *0.6033* | 0.075 |
| 2I | labeled in >1 location | 6.76 | 0.0137 | 0.166 | 3.04 | 0.0304 | 0.263 |
| 2J | labeled head only | 6.02 | 0.0194 | 0.151 | 0.53 | *0.7162* | 0.058 |
| 2K | labeled head and neck | 7.28 | 0.0108 | 0.176 | 2.62 | *0.0520* | 0.236 |
| 3C | percent spines labeled | 18.53 | 0.0001 | 0.353 | 0.80 | *0.5306* | 0.087 |
| 3D | percent spine heads labeled | 16.85 | 0.0002 | 0.319 | 1.44 | *0.2408* | 0.138 |
| 3D | percent spine necks labeled | 8.37 | 0.0064 | 0.189 | 2.09 | *0.1018* | 0.189 |
| 4B | labeled, PSD < 0.1 $\mu\text{m}^2$ | 8.41 | 0.0065 | 0.198 | 0.22 | *0.9228* | 0.026 |
| 4B | labeled, PSD > 0.2 $\mu\text{m}^2$ | 8.54 | 0.0061 | 0.201 | 1.43 | *0.2437* | 0.144 |
| 4D | unlabeled, PSD < 0.15 $\mu\text{m}^2$ | 4.47 | 0.0420 | 0.116 | 1.71 | *0.1703* | 0.168 |
| 4F | labeled head, PSD < 0.1 $\mu\text{m}^2$ | 5.37 | 0.0266 | 0.136 | 0.49 | *0.7438* | 0.054 |
| 4F | labeled head, PSD > 0.2 $\mu\text{m}^2$ | 8.54 | 0.0061 | 0.201 | 1.43 | *0.2437* | 0.144 |
| 5D | labeled, with spine apparatus | 10.93 | 0.0022 | 0.243 | 1.60 | *0.1979* | 0.158 |
| 5D | unlabeled, with spine apparatus | 5.37 | 0.0266 | 0.137 | 2.38 | *0.0710* | 0.219 |
| 5D | labeled, without spine apparatus | 4.78 | 0.0357 | 0.123 | 0.55 | *0.6997* | 0.061 |
| 5D | unlabeled, without spine apparatus | 4.19 | 0.0485 | 0.110 | 1.27 | *0.3026* | 0.130 |
| 5F | head labeled, with spine apparatus | 10.93 | 0.0022 | 0.243 | 2.06 | *0.1083* | 0.195 |
| 5F | neck labeled, with spine apparatus | 13.17 | 0.0009 | 0.279 | 6.32 | 0.0006 | 0.427 |
| 5F | base labeled, with spine apparatus | 5.39 | 0.0264 | 0.137 | 2.08 | *0.1056* | 0.196 |
| 5F | head labeled, without spine apparatus | 4.37 | 0.0442 | 0.114 | 1.61 | *0.1954* | 0.159 |
| 5I | labeled, with spine apparatus, PSD < 0.15 $\mu\text{m}^2$ | 4.40 | 0.0435 | 0.114 | 0.94 | *0.4548* | 0.099 |
| 5F | labeled, with spine apparatus, PSD > 0.2 $\mu\text{m}^2$ | 8.54 | 0.0061 | 0.201 | 1.43 | *0.2437* | 0.144 |

## Results

To investigate dendritic eIF distribution during memory consolidation, we used aversive Pavlovian conditioning (also known as threat or fear conditioning), an extensively studied behavioral paradigm which produces robust learning in a single training session (LeDoux, 2000; Maren, 2001). Formation of the long-term memory is impaired by intra-LA infusion of general protein synthesis inhibitors (Schafe and LeDoux, 2000; Maren et al., 2003) or 4EGI-1 (Hoeffer et al., 2011) immediately after training. The memory is resistant to a protein synthesis inhibitor infused after 6 hours, confirming the importance of translation in the first few hours (Schafe and LeDoux, 2000). The temporal dynamics of learning-induced translation in the LA have not been worked out in detail, but studies of memory formation in hippocampus consistently report a protein synthesis-dependent phase lasting less than three hours after learning, followed sometimes by a second phase whose timing varies between learning paradigms (Bourtchouladze et al., 1998; Quevedo et al., 1999). A variety of signaling pathways induced by a single trial of avoidance learning peak within 30 to 90 minutes across several brain areas but diverge thereafter, further supporting the importance of molecular changes during the first hours (Izquierdo et al., 2006). To capture eIF distribution associated with this period of active memory formation, we trained animals with five tone-shock pairings and collected tissue 1 hour after the first pairing, which was 32.5 minutes after the last pairing. In previous work, we found an increase in polyribosomes in spines after paired training relative to training with unpaired shocks and tones or or control exposure to the training box (Ostroff et al., 2010, 2017). Polyribosome distribution did not differ between unpaired training and box exposure, but unpaired training did produce inhibitory learning and reduced synapse size. Because ssTEM entails a large investment in each subject, we chose to use box exposure as the behavioral control for paired training, as it does not induce learning. The experimental workflow is shown in Figure 1A. Freezing was scored during training to confirm acquisition of the tone-shock association in the paired group (Figure 1B).

**Figure 1.**
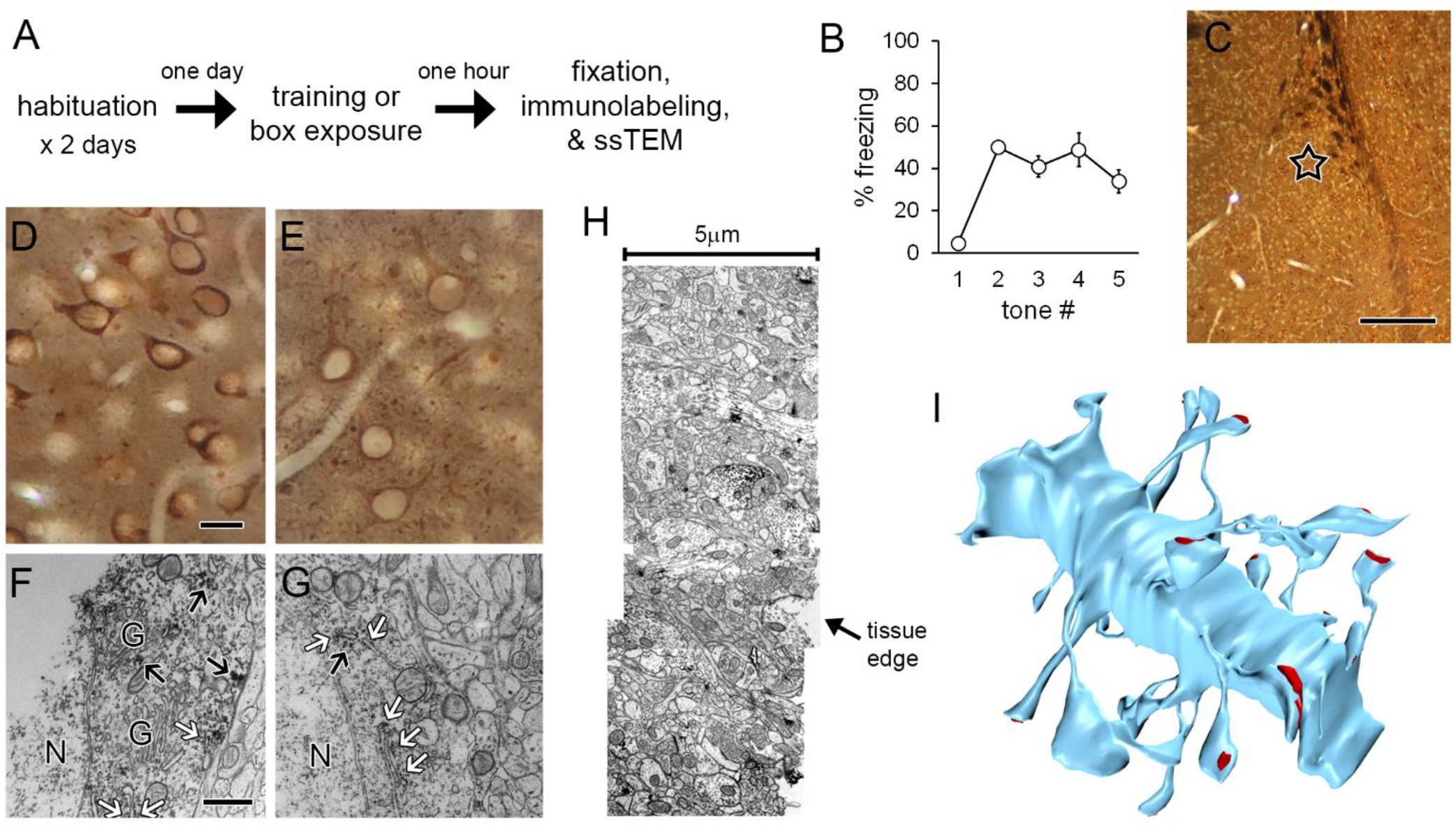
Methods. A) Experimental workflow. B) Freezing to tones during the training session by the rats used for ssTEM. C) Lateral amygdala section immunolabeled for eIF4E and embedded for EM, showing location of sampling for ssTEM (star). D-E) Brightfield image of LA cells labeled for eIF4E (E) and eIF4G1 (E) in EM sample blocks. F-G) Electron micrographs of cell bodies showing labeling for eIF4E (F) and eIF4G1 (G) on rough endoplasmic reticulum (white arrows) and on cytoplasmic polyribosomes (black arrows). H) Montage of three sections along edge of tissue labeled for eIF4E. I) Dendrite from eIF4E labeled tissue reconstructed by ssTEM. Synapses shown in red. Scale = 250 mm in C; 20 mm in D-E; 500 nm in F-G.

To determine whether eIF4E and eIF4G are present in LA dendrites and spines, we performed pre-embedding immunohistochemistry for these proteins on alternating sections before processing the tissue for ssTEM. A region of the dorsolateral subdivision of the LA at a mid-caudal level was chosen for ssTEM (Figure 1C). For optimal sensitivity of immunolabeling, we used a chromogenic detection method and only used one antibody on each tissue section. Although this approach does not reveal colocalization, it maximizes the likelihood of detecting labeled spines, as the antibodies and labeling reagents do not interfere with each other.

For both eIF4E and eIF4G1, polyclonal antibodies raised against the C-terminal region were used. The site of interaction between eIF4E and eIF4G1 is not near the C-terminal of either molecule (Mader et al., 1995; Morino et al., 2000; Grüner et al., 2016), and it is therefore possible that the antibodies recognize both the unbound proteins and the eIF4F complex. The 4E binding proteins (4E-BPs), which are major repressors of eIF4E activity, bind at the eIF4G binding site and similarly should not directly block the C-terminus (Marcotrigiano et al., 1999; Tomoo et al., 2005). The C-terminus of eIF4E is part of the cap-binding pocket and is deflected, but not occluded, by binding to mRNA (Marcotrigiano et al., 1997; Tomoo et al., 2002). The C-terminus of eIF4G, meanwhile, does not bind the protein’s other major binding partner, eIF4A, and appears to be modulatory (Morino et al., 2000).

At the light level, labeling for both initiation factors was seen in the cytoplasm of the cell bodies, proximal dendrites, and processes throughout the neuropil, but not in the nuclei (Figure 1D-E). At the EM level, immunolabel in cell bodies was found associated with rough endoplasmic reticulum and with free polyribosomes as well as sporadically in the cytoplasm, but was conspicuously absent from the Golgi apparatus (Figure 1F-G). This staining pattern is consistent with expected subcellular sites of translation. Because the antibodies only penetrated a few microns into fixed tissue, analysis was restricted to the region closest to the vibratome edge. To maximize our usable image volumes, three adjacent fields along the tissue edge were imaged and montaged (Figure 1H). Spiny dendrites and their synapses were reconstructed in 3D (Figure 1I) and the distribution of immunolabel was quantified.

### Learning-induced upregulation of eIF4E in dendritic spines

Immunolabeling for both eIF4E and eIF4G1 was visible in dendritic spine heads (Figure 2A-B), necks (Figure 2C), and bases (Figure 2D). Dendritic shafts Quantification of the frequency of labeled and unlabeled spines along dendritic segments in the control group revealed that a majority of spines were labeled for eIF4G1, but not for eIF4E. In the trained group, there were more spines labeled for eIF4E and fewer unlabeled spines, but no changes in eIF4G labeling (Figure 2E). The mean frequencies of spines with eIF4G1 labeling in both groups were nearly identical to the frequency of spines with eIF4E labeling in the trained group. Although this is intriguing to note, direct comparison between the two patterns could be misleading because the detection sensitivity of the antibodies may differ. The frequency of labeled and unlabeled spines was not correlated for either antibody in either condition (Figure 2F-G), suggesting that the presence of labeling is not directly regulated at the level of the dendritic segment. When the presence of labeling was broken down by location in spines, more spines with eIF4E labeling in the head and neck were found in the trained group, but there were no effects on eIF4G1 labeling (Figure 2H). In the trained group the most frequent location for eIF4E label was the spine head, while spine bases were more frequently labeled for eIF4G1.

**Figure 2.**
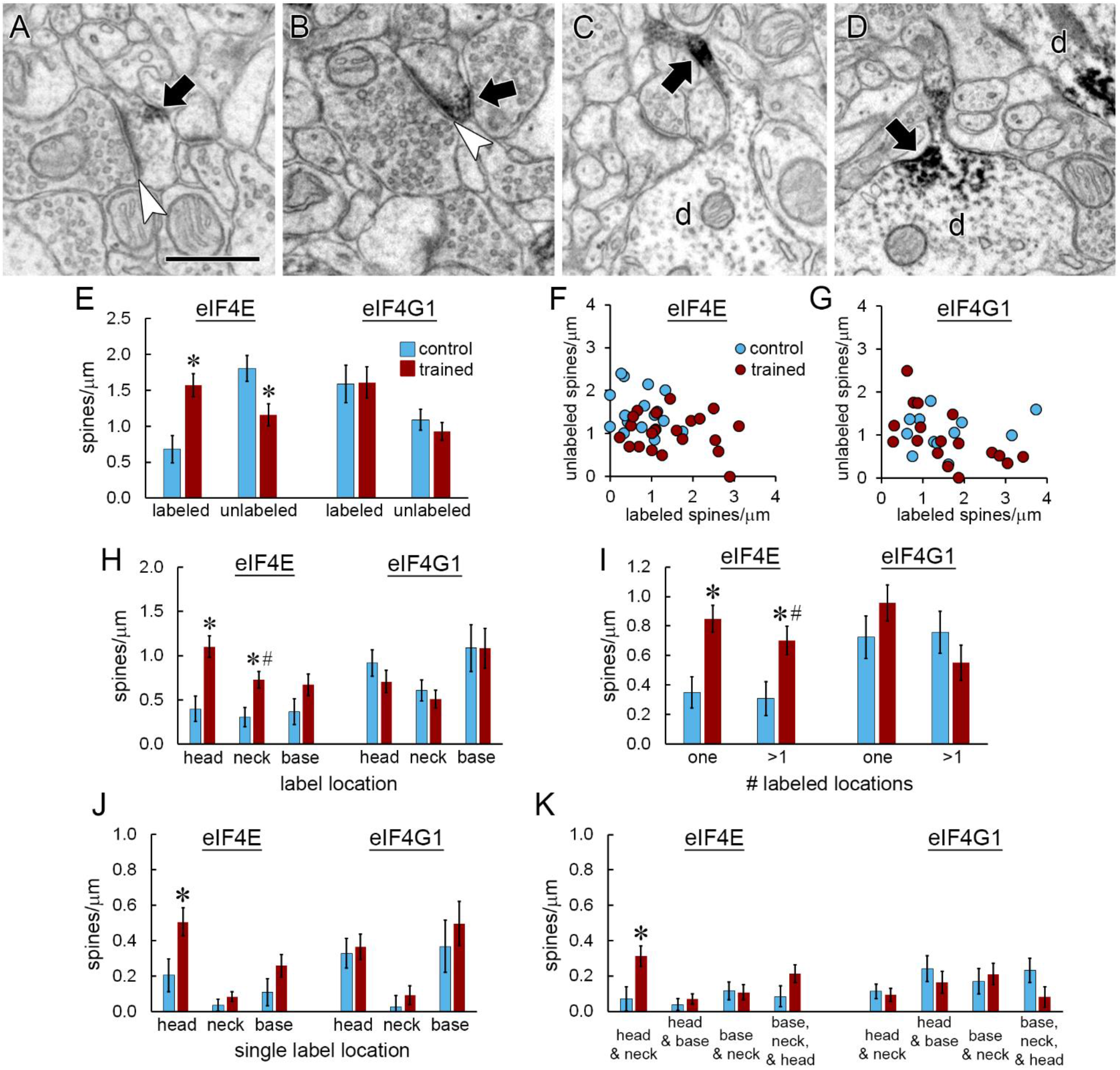
Distribution of immunolabel for eIF4E and eIF4G1. A-B) Immunolabel (arrows) for eIF4E (A) and eIF4G1 (B) in the heads of dendritic spines forming asymmetric synapses (arrowheads). C-D) Dendrites (d) with immunolabel (arrows) for eIF4E in a spine neck (C) and a spine base (D). Another dendrite (d) containing label in its shaft is visible in the upper right corner of (D). Scale = 500nm. E) The frequency of spines labeled for eIF4E, but not eIF4G, increased after learning, with a corresponding decrease in unlabeled spines. F-G) There were no correlations between the frequency of labeled and unlabeled spines for eIF4E (F) or eIF4G1 (G). H) The frequency of spine heads and necks labeled for eIF4E, but not eIF4G1, increased after learning. I) The frequency of spines with eIF4E label in one or multiple locations increased with learning. J) Among spines with only one labeled location, only eIF4E-labeled spine heads increased. K) Among spines with multiple labeled locations, spines with eIF4E in the head and neck were increased. * p < 0.05

To examine how labeling was distributed within spines, we classified labeled spines by the number of labeled locations. Labeling for eIF4E and eIF4G1 was commonly seen in multiple locations within a spine, with similar numbers of spines having label in single and multiple locations. More spines were found with both labeling patterns for eIF4E in the trained group (Figure 2I). Of the spines with label in a single location, the increase in eIF4E-labeled spines was specific to the head location (Figure 2J). Very few spines had labeling for either eIF4E or eIF4G1 in the neck only, suggesting that labeling in the neck reflects diffusion towards or away from accumulation in the base or head. Of spines with multiple labeled locations, only those with both head and neck label were significantly increased in the trained group (Figure 2K). Overall, these analyses show upregulation of spines with eIF4E labeling in the head, and sometimes also in the neck, during memory consolidation. In contrast, eIF4G1 labeling in spines is unaffected by training.

### Distribution of eIF2α in dendritic spines

In addition to eIF4E interaction, the second major target of translational control mechanisms is eIF2α, whose regulation has been implicated in amygdala-based memory (Trinh and Klann, 2013; Jian et al., 2014). To investigate whether the distribution of eIF2α in spines resembles that of eIF4E, we repeated the learning experiment on a new cohort of animals and immunolabeled the tissue for eIF2α. The antibody we used was raised against the full-length protein, so in theory it could recognize eIF2α whether free or in the preinitiation complex. Qualitatively, labeling for eIF2α resembled that of eIF4E and eIF4G1, with reaction product distributed throughout the cell bodies, proximal dendrites, and neuropil processes, but largely sparing the nucleus (Figure 3A). Like the other initiation factors, eIF2α labeling was found in dendritic spine heads (Figure 3B), as well as necks and bases. Because of the tissue preparation conditions required for efficient labeling of eIF2α, the morphology of this material was relatively poorer and the yield of dendrites suitable for reconstruction was lower than in the eIF4E-eIF4G1 experiment. Many spines could not be unequivocally reconstructed along the dendrites thereby rendering absolute frequency measurements inaccurate, so we limited analysis of this material to percentages of labeled spines. Whereas the percentage of spines labeled for eIF4E increased with training, the percentage of spines labeled for eIF2α, like eIF4G1, was unchanged (Figure 3B). At the sub-spine level, the percentage of spine heads and necks with eIF4E label was higher in the trained group (Figure 3D), but there was no effect of training in any location for eIF4G1 (Figure 3E) or eIF2α (Figure 3F).

**Figure 3.**
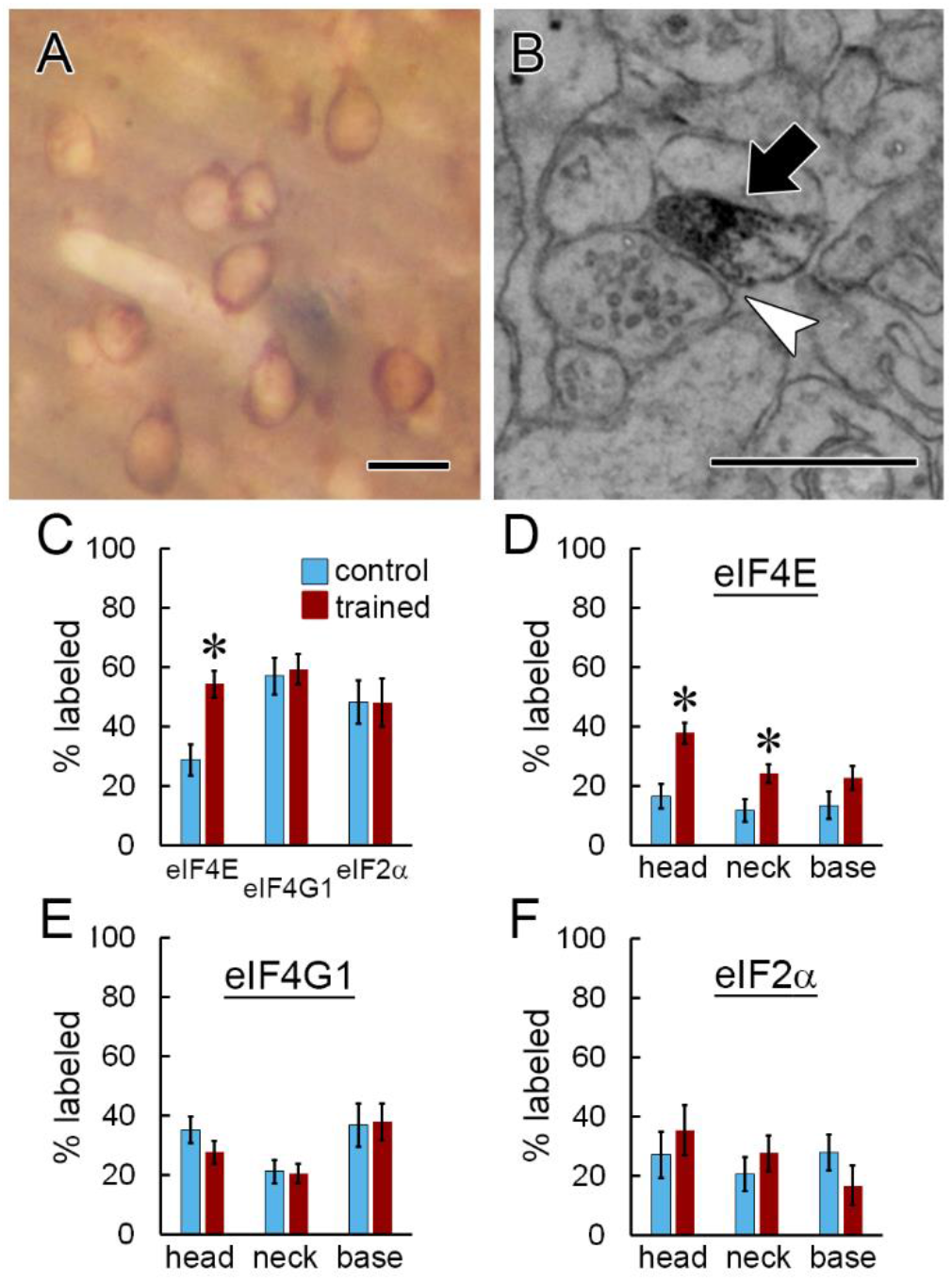
Comparison between eIF4E, eIF4G1, and eIF2α distribution. A) Brightfield image of eIF2α immunolabel in LA tissue prepared for EM. B) EM image of eIF2α immunolabeling in a dendritic spine head (arrow) forming an asymmetric synapse (arrowhead). Scale = 500nm. C) A higher percentage of dendritic spines are labeled for eIF4E, but not eIF4G1 or eIF2α, after learning. D-F) A higher percentage of dendritic spines heads and necks contain eIF4E (D), but not eIF4G1 (E) or eIF2α (F), after learning. * p < 0.05 Scale = 20mm in A; 500 nm in B.

### Accumulation of eIF4E label in large and small spine heads

The spine enlargement that occurs with synapse strengthening is believed to be a substrate for long-term memories (Bourne and Harris, 2007; Kasai and Fukuda, 2010), and translation is necessary for plasticity-associated spine enlargement in the rat hippocampus (Fifkova et al., 1982; Tanaka et al., 2008). Proliferation of small spines is associated with eIF4E manipulation; small spine outgrowth was observed in the LA when 4EGI-1 was infused after training (Ostroff et al., 2017), but was also seen in the prefrontal cortex of mice overexpressing eIF4E (Santini et al., 2013). To determine whether eIF distribution differed between spines of different sizes, we measured the area of the post-synaptic density (PSD), which is correlated with spine volume and surface area in the LA (Ostroff et al., 2010). PSD area was quantified in two dimensions for all spines by directly measuring the area of PSDs cut *en face* or by measuring the length of cross-sectioned PSDs across serial sections as shown in Figure 4A. Synapse size ranged from 0.003 - 0.849 μm^2^ and did not follow a normal distribution, with nearly 90% of the synapses in the dataset smaller than 0.1 μm^2^. To determine how labeling for eIF4E and eIF4G1 were distributed across the range of spine sizes, spine frequency was binned. There were more eIF4E-labeled spines in the trained group at both the smaller (0.05 - 0.1 μm^2^) and largest (> 0.2 μm^2^) ends of the range (Figure 4B), and fewer unlabeled spines in the intermediate (0.1 - 0.15 μm^2^) range (Figure 4C). Spines with eIF4G1 labeling spanned the full range of synapse sizes, but there were no differences between the training groups (Figure 4D). As seen for eIF4E, large spines lacking eIF4G1 label were very scarce (Figure 4E). When eIF4E-labeled spines were analyzed by size and label location, the trained group had more labeled spine heads in the 0.05 - 0.1 μm^2^ and >0.2 μm^2^ bins (Figure 4F), but no size-specific effects were seen in the neck (Figure 4G) or base (Figure 4H) locations.

**Figure 4.**
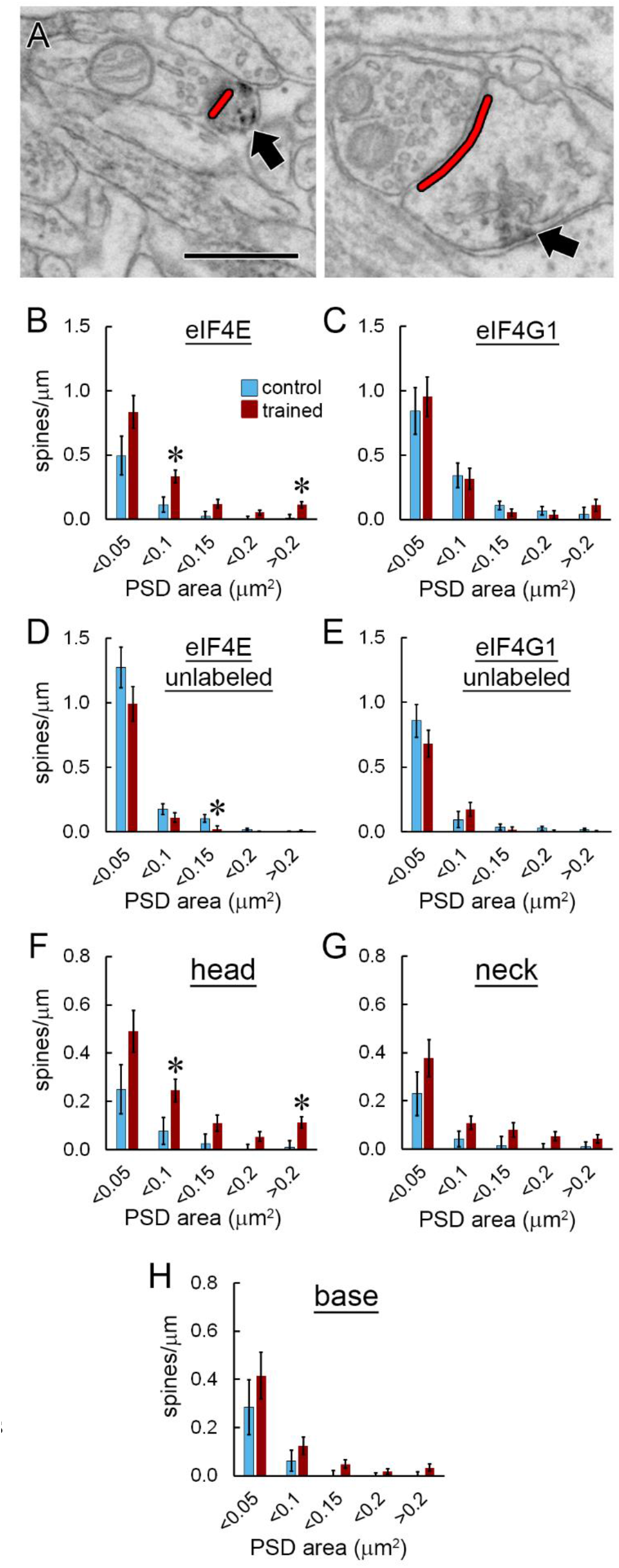
Synapse size. A) EMs showing measurement of postsynaptic density length (red) on spine heads containing label for eIF4E (arrows). Scale = 500 nm. B-E) Spine frequency binned by synapse size for spines with (B) or without (C) eIF4E labeling and with (D) or without (E) eIF4G1 labeling. F-H) Frequency of spines binned by synapse size with eIF4E labeling in the head (F), neck (G), or base (H). * p < 0.05

### Differential distribution of labeling in spines with and without a spine apparatus

The spine apparatus is a membranous organelle found in the largest dendritic spines (Westrum et al., 1980; Spacek, 1985; Ostroff et al., 2010). It is a specialization of the continuous smooth endoplasmic reticulum (sER) network that extends throughout the dendritic arbor, and consists of membranous calcium-containing cisterns interleaved with dense actin plates (Fifkova et al., 1983; Deller et al., 2000; Capani et al., 2001). The functional activities of the spine apparatus have not been determined, but it contains Golgi markers (Pierce et al., 2001) and thus could have a role in post-translational processing as well as calcium signaling. Immunolabeling for eIFs was found in spines with and without a spine apparatus (Figure 5A-C). The overall effect of training on spines with eIF4E labeling was similar in spines with and without a spine apparatus: the trained group had more labeled and fewer unlabeled spines of both types (Figure 5D). There was no effect of training on eIF4G1 distribution with respect to the spine apparatus (Figure 5E). Within spines, the trained group had more labeling for eIF4E in the heads, necks, and bases of spines with a spine apparatus, but only in the heads of spines without a spine apparatus (Figure 5F). Again, there were no training effects on eIF4G1 labeling (Figure 5G).

**Figure 5.**
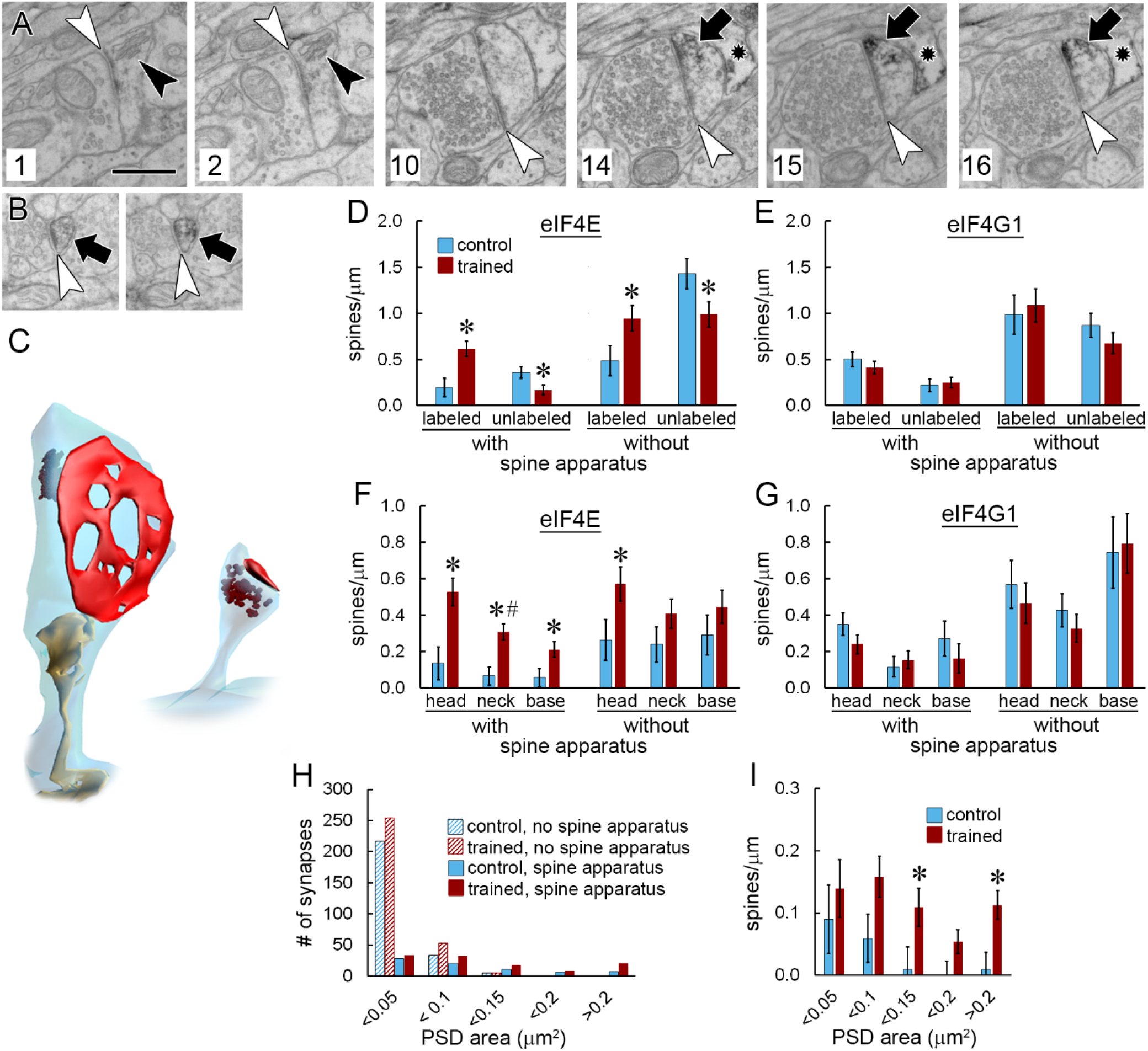
Spine apparatus. A) Six non-consecutive serial EM images through a large spine forming an asymmetric synapse (white arrowheads) and containing a spine apparatus (black arrowheads) and immunolabel for eIF4E (arrows). Section numbers are shown at lower left. An astrocytic process (star) containing immunolabel is visible in sections 14-16. Scale = 500nm. B) Two consecutive serial EM images of a spine without a spine apparatus labeled for eIF4E. C) 3D reconstructions of the spines shown in (A) and (B) with spine apparatus in yellow, synapse in red, and label in brown. D-E) Frequency of spines with and without eIF4E (D) or eIF4G1 (E) labeling broken down by the presence of a spine apparatus. F-G) Spine frequency by label location and presence of a spine apparatus for eIF4E (F) and eIF4G1 (G). H) Histogram showing the synapse size distribution for all spines in the dataset by training group and presence of a spine apparatus. I) Frequency of spines with a spine apparatus and eIF4E immunolabel, binned by synapse size. * p < 0.05; # significant interaction with subject.

Spines that contain a spine apparatus are several-fold larger on average than other spines (Ostroff et al., 2010); across all spines in the analyzed dataset, spines without a spine apparatus had a mean synapse size of 0.034 ± 0.003 μm^2^, compared to 0.128 ± 0.006 μm^2^ for spines with a spine apparatus. We find that nearly all spines with synapses >0.1 μm^2^ contain a spine apparatus, making it a universal feature of the largest spines. The spine apparatus is not exclusively found in large spines, however, but is present across the entire range of synapse sizes (Figure 5H). When spines with a spine apparatus were binned by synapse size, the trained group had more eIF4E-labeled spines only in the largest (>0.2 μm^2^) and intermediate (0.1 - 0.15 μm^2^) bins (Figure 5I). Among small spines, therefore, training only affects eIF4E labeling in those without a spine apparatus.

### eIF labeling in dendritic shafts

Immunolabel for eIFs was present in the majority of dendritic shafts. As in dendritic spines, the DAB reaction product in shafts was not found in the form of isolated puncta, but typically consisted of irregular groupings of dark puncta and patches of speckling spread through swaths of cytoplasm (Figure 6A). The diffuse, amorphous nature of the label does not allow quantification of the number of antibody molecules. We instead performed a semi-quantitative evaluation of label distribution in the shafts by measuring the volume of cytoplasm that contained labeling as a percentage of each dendrite’s cylindrical shaft volume. Labeling for eIF4E was present throughout an average of 7.1% of the shaft cytoplasm (range: 0 - 32.4%), and eIF4G1 labeling was in 30.7% (range: 0 - 100%). There were no differences between training groups (Figure 6B). Although this is not a quantitative analysis of protein levels, the range of values suggests that elFs are non-uniformly distributed in dendritic shafts and that eIF4G1 has a more extensive presence than eIF4E. Plotting the labeled shaft volume against dendrite diameter showed no clear relationship (6C-D). The frequency of unlabeled spines likewise appears unrelated to the extent of shaft labeling (Figure 6E-F), but the frequency of labeled spines looks to be higher in the most heavily labeled dendrites relative to the least (Figure 6G-H).

**Figure 6.**
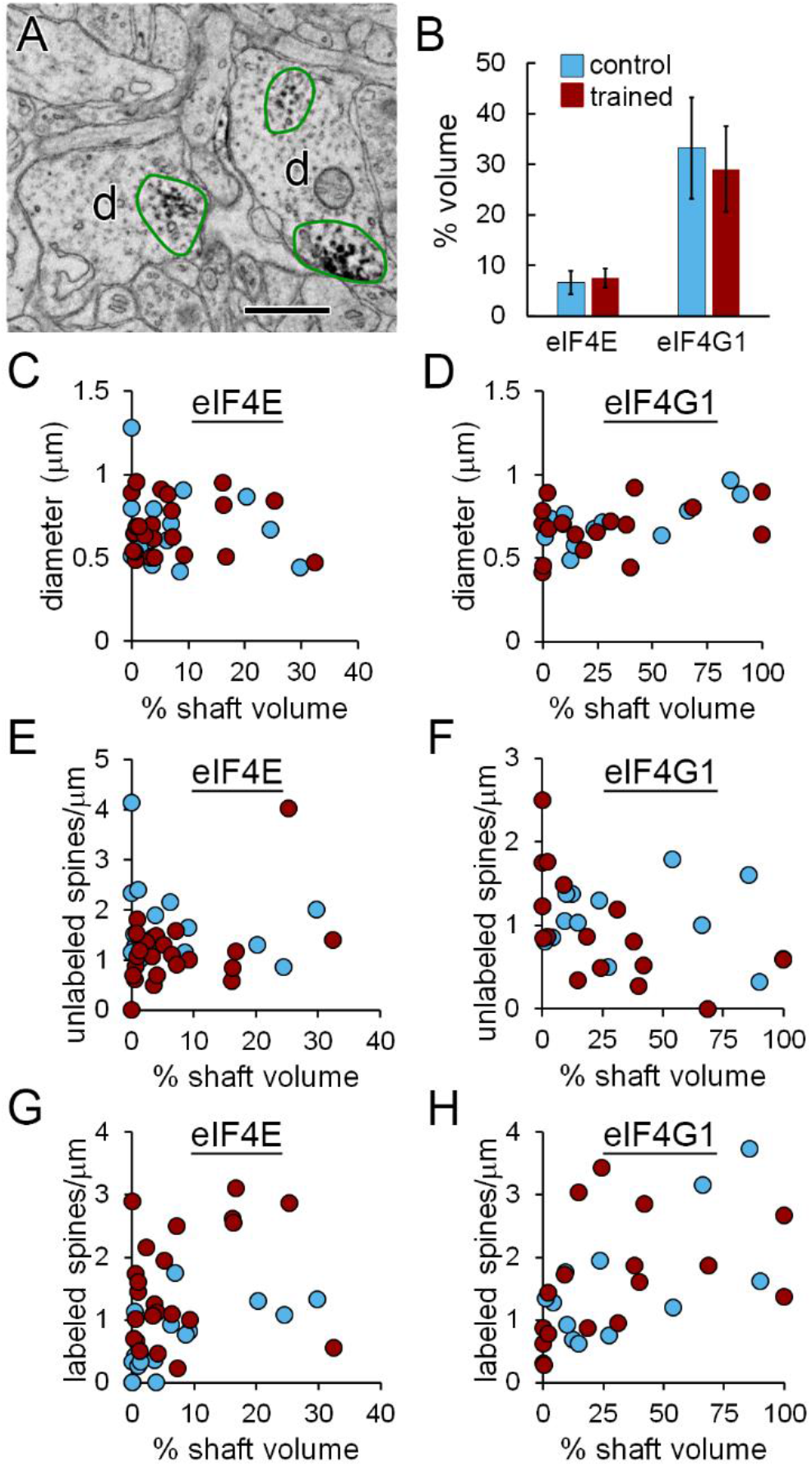
Labeling in dendritic shafts. A) Two dendrites (d) with patches of cytoplasm containing immunolabeling for eIF4E outlined in green. B) Average percent of dendritic shaft volume containing immunolabel for eIF4E or eIF4G1. C-D) Plots of percent shaft volume with label for eIF4E (C) and eIF4G1 (D) versus diameter for each dendrite. E-H) Plots of percent shaft volume with label for eIF4E (E,G) and eIF4G1 (F,H) versus frequency of unlabeled spines (E-F) and labeled spines (G-H).

### Comparison of eIF labeling with polyribosome distribution in spines

If polyribosomes in spines are locally assembled via cap-dependent initiation, we would expect some correspondence between the spine distribution of eIFs and polyribosomes. Due to the nature of our labeling procedure, we were unable to reliably identify polyribosomes in dendritic spines in this study. To preserve contrast between the label and the tissue, we omitted the usual counterstaining that enhances polyribosomes. A few polyribosomes were visible amid deposits of immunolabel in spines (Figures 1A, 5A-16, and 5B-1 for example), as well as in dendrites and especially in cell bodies, where they are highly abundant, but none were observed in unlabeled dendritic spines. Because of the lack of counterstaining, we cannot conclude that these spines lack polyribosomes. Likewise, the absence of polyribosomes in labeled spines could not be determined because the immunolabeling was often dense enough to occlude structures in the cytoplasm.

The training protocol, timing of tissue collection, and analysis methods used here were identical to those used in our ssTEM study of the effects of 4EGI-1 on polyribosomes (Ostroff et al., 2017), so we compared the frequencies of spine polyribosomes from that study (Figure 7A) with eIF labeling to put the datasets in context. Polyribosomes were found in approximately 30% of dendritic spines in the vehicle-infused control group and approximately half of spines in the vehicle-infused trained group, similar to the percentages of spines labeled for eIF4E (Figure 7B). In both studies, effects differed by sub-spine location and the presence of a spine apparatus, so we compared these measures as well. In the heads of spines with a spine apparatus, both eIF4E labeling and polyribosomes were more frequent in the trained groups, and 4EGI-1 blocked the effect of training but did not reduce the control level of polyribosomes (Figure 7C). Both eIF4E and polyribosomes were also upregulated in the heads of spines without a spine apparatus, but in this case both drug groups did have fewer polyribosomes (Figure 7D). The spine base and neck locations did not differ, so spines with labeling or polyribosomes in the base, neck, or both were pooled. There was an effect of training on eIF4E labeling, but not polyribosomes, in the base/neck location of spines with a spine apparatus (Figure 7E). In the bases and necks of spines without a spine apparatus, there were more polyribosomes in the trained groups with no effect of 4EGI-1, and no significant effect on eIF4E (Figure 7F).

**Figure 7.**
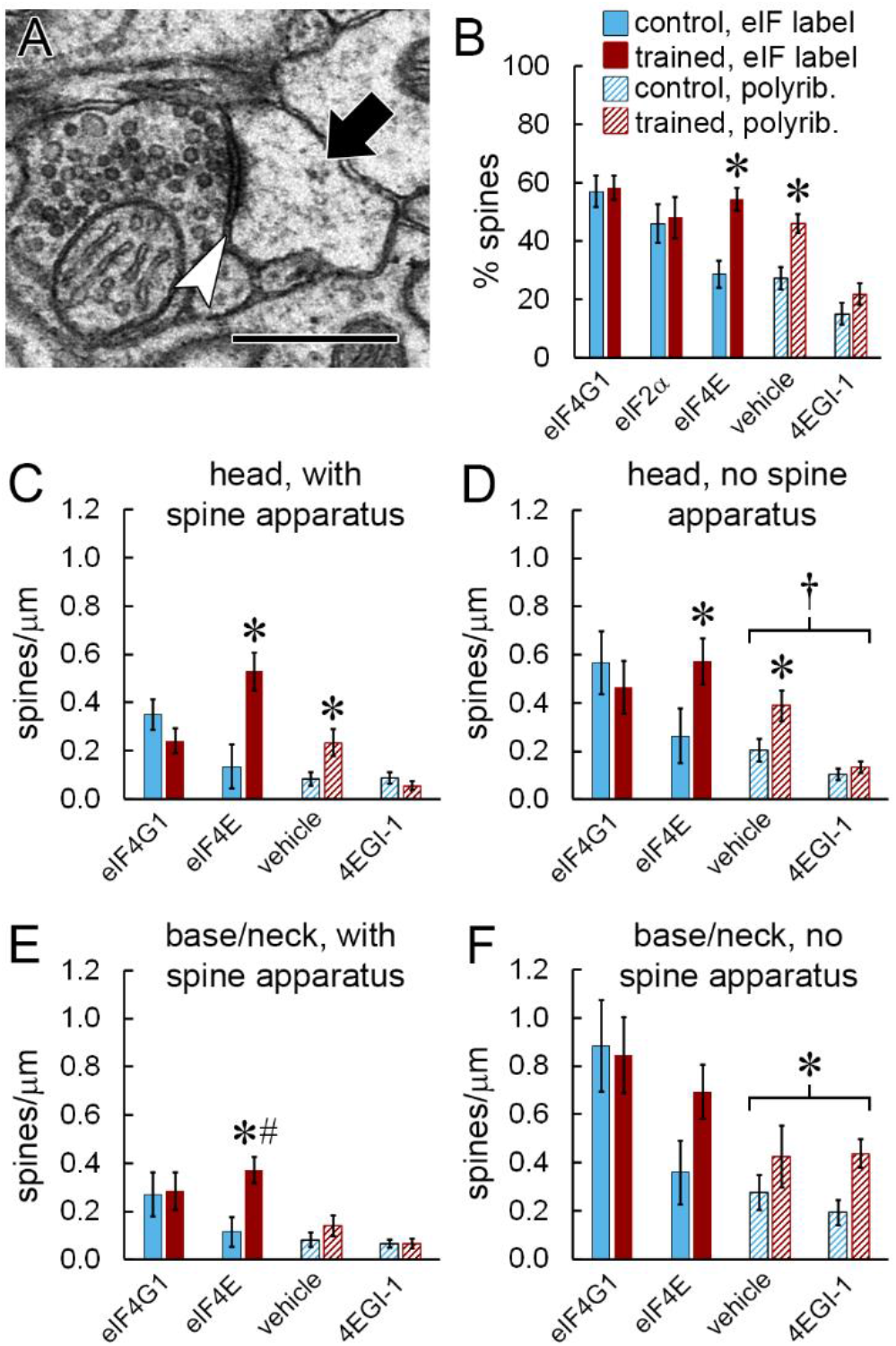
Comparison with polyribosome distribution. A) EM of a polyribosome (arrow) in the head of a dendritic spine forming a synapse (arrowhead). Scale = 500 nm. B) Percentage of all spines labeled for eIFs or containing polyribosomes. C-F) Effects of learning on the frequency of spines with immunolabel for eIF4G1 or eIF4E, or with polyribosomes after infusion of vehicle or 4EGI-1. Data are broken down by location of label in the spine head (C,D) or base/neck (E,F) and the presence (C,E) or absence (D,F) of a spine apparatus. Labeling data in (C) and (D) replotted from Figure 5F-G, and all polyribosome data replotted from Ostroff et al. (2017). * p < 0.05; # significant interaction with subject; † effect of 4EGI-1

The data shown in Figure 7 are summarized in Figure 8. The effect of learning on eIF4E distribution was similar to the effect on cap-dependent (4EGI-1-sensitive) but not cap-independent (4EGI-1-insensitive) polyribosomes: eIF4E and cap-dependent polyribosomes accumulated in the heads of dendritic spines, while cap-independent polyribosomes, but not eIF4E, accumulated in the bases and necks of spines lacking a spine apparatus. Although the other eIFs were distributed throughout dendrites and spines, the selective upregulation of eIF4E in locations where cap-dependent polyribosomes accumulate is consistent with eIF4E serving as the rate limiting component of initiation. Overall, our data indicate that the machinery for translation initiation is localized near individual synapses, and that initiation may be regulated in a highly spatially-specific manner during memory consolidation via availability of eIF4E.

**Figure 8.**
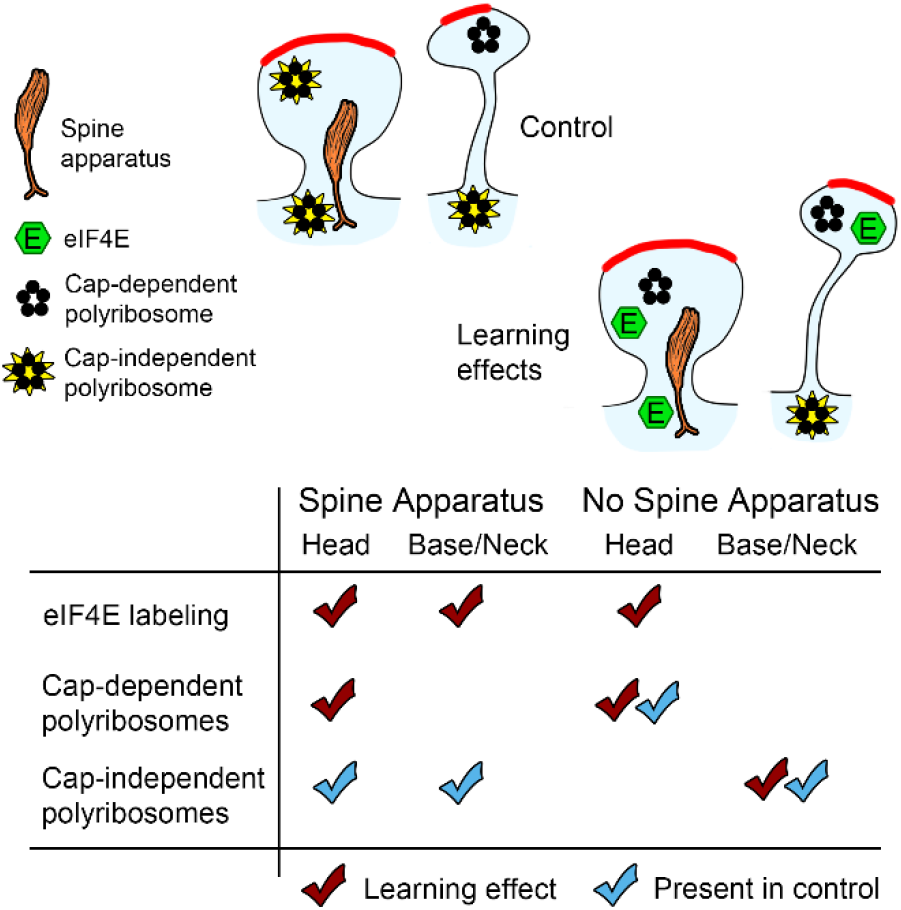
Summary of learning effects on eIF4E labeling, cap-dependent (4EGI-1 sensitive) and cap-independent (4EGI-1 insensitive) polyribosomes. Effects on eIF4E from this study, effects on polyribosomes from Ostroff et al. (2017).

## Discussion

Our results indicate that three different eIFs localize to dendritic spines, supporting the hypothesis that *de novo* protein synthesis can be regulated at the level of individual synapses. Targeting specific proteins to individual synapses is presumed to be essential for maintaining the fidelity of memory networks. Local translation is an obvious candidate mechanism, as it can provide proteins rapidly at a distance from the nucleus, potentially with very high spatiotemporal precision. Because mRNA trafficking and translation are heavily regulated and multiple signaling pathways must converge in a single location, translation effectively serves as a coincidence detector. Synapse-specific translation regulation could be achieved by combinations of selective spatial targeting and biochemical modulation of mRNA, translation machinery, or both. A variety of dendritic mRNA transport mechanisms have been identified and appear to be selective for specific transcripts (Miller et al., 2002; Buxbaum et al., 2015; Nakayama et al., 2017; Roy et al., 2020), and mRNA can remain dormant in dendrites until it is unmasked by synaptic activity (Hutten et al., 2014). Certain mRNAs have been observed in spine heads while others are confined to the base, suggesting that mRNA targeting can be spatially precise at the spine level (Tiruchinapalli et al., 2003; Dynes and Steward, 2012). Downstream of mRNA trafficking, translation machinery offers another potential substrate for synapse-level regulation, and such a role could explain why intellectual functions such as learning are uniquely sensitive to disruption of translation regulation (Kelleher and Bear, 2008; Darnell, 2011; Kapur et al., 2017). Our data are consistent with earlier reports of eIF4E labeling in the spines of cultured neurons (Smart et al., 2003) and cortex (Asaki et al., 2003) and furthermore suggest that eIF4E, considered the key control point in translation (Amorim et al., 2018), is specifically regulated in spines during memory consolidation.

Biochemical regulation of eIFs alone could allow nuanced translational control in spines. For example, as multiple different signaling pathways and effector molecules separately target eIF4E and eIF2α (Trinh and Klann, 2013; Amorim et al., 2018), cap-dependent translation in a spine could require concurrent activation of potentially orthogonal triggers. Two major signaling molecules with well-established roles in multiple forms of synaptic plasticity, memory, and neurological disease are key regulators of eIF4E: the mechanistic target of rapamycin complex 1 (mTORC1) and mitogen-activated kinase (MAPK) pathways (Hoeffer and Klann, 2010; Kelleher et al., 2004). Both pathways are downstream of synaptic activity, but they target eIF4E differently. By phosphorylating 4E-BPs, mTORC1 decreases their affinity for eIF4E and thereby facilitates translation, while MAPK activates MAPK-interacting kinases (MNKs) which phosphorylate eIF4E itself, also facilitating translation (Marcotrigiano et al., 1999; Pyronnet et al., 1999; Shveygert et al., 2010; Amorim et al., 2018). MNK is recruited to phosphorylate eIF4E by eIF4G (Pyronnet et al., 1999; Shveygert et al., 2010), meaning that the ability of MAPK to trigger initiation may depend on prior binding of eIF4E to eIF4G. The eIF4G binding site of eIF4E can also be occluded by a complex of the fragile X mental retardation protein (FMRP) and the cytoplasmic FMRP interacting protein (CYFIP1), which are released from eIF4E by MNK activity (Genheden et al., 2015). Regulation of eIF2α is less studied than that of eIF4E, but its activity is modulated by kinases whose activity is necessary for normal learning and cognitive function, suggesting that it too could have a role in local translation (Costa-Mattioli et al., 2007; Trinh and Klann, 2013).

Because we used eIF antibodies that were not specific to phosphorylation or complexing with major regulators, we were able to observe the dynamics of eIF localization separately from biochemical regulation. Fewer than 60% of spines contained labeling for any eIF, and only a third of spines were labeled for eIF4E in the control condition. Similar proportions of spines were labeled for eIF4G1 and eIF2α regardless of training, but fewer were labeled for eIF4E in the control group. The proportion of spines with eIF4E labeling increased in the trained group, suggesting a scenario in which some spines are primed for translation by the presence of eIF4G1 and eIF2α, but initiation does not occur until synaptic activity triggers delivery of eIF4E. This is consistent with the canonical view that eIF4E is the rate limiting factor in initiation, but without multiplexed labeling we do not know how often all three eIFs are present in the same spines. The four-fold difference between the spatial extent of eIF4G1 versus eIF4E labeling in the dendritic shafts also argues for limited eIF4E availability as a regulatory mechanism. The possibility of differences in sensitivity between the antibodies makes direct comparisons difficult, however.In addition, it is possible that our antibody was less efficient at binding to eIF4E in complex with its repressor proteins, and the increased labeling reflects activation of pre-existing eIF4E. Interestingly, a quantitative proteomic analysis of HeLa cells found similar protein copy numbers of eIF4G1 and eIF2α, which were almost double that of eIF4E (Kulak et al., 2014). Copy number data are not available for neurons or their compartments, but could shed light on the relative role of local copy number versus biochemical mechanisms in translation regulation.

A particularly intriguing aspect of our data is the correspondence between eIF4E distribution and the distribution of polyribosomes in our earlier study (Ostroff et al., 2017). Although this supports the general model of eIF4E as rate-limiting, it may also reflect something more specific. Polyribosomes are traditionally assumed to represent a default mode of translation, with multiple ribosomes appearing in a rosette shape on an mRNA circularized by the interaction of the cap-eIF4E-eIF4G complex with the poly-A binding protein (Vicens et al., 2018). Recent evidence indicates that things are not so simple. First, polyribosomes are not the only sites of cytoplasmic translation: two studies have reported that a substantial amount of translation occurs on monosomes, with one showing that in neurons dendritic translation is especially biased towards monosomes (Heyer and Moore, 2016; Biever et al., 2020). Second, polyribosomes may not be actively translating; stalled polyribosomes may be transported in dendrites for later reactivation, and live imaging of cultured neurons found that dendritic polyribosomes are nearly completely stalled (Richter and Coller, 2015; Langille et al., 2019). Third, although eIF4E-dependent initiation is considered the default translation pathway, there is growing evidence that a significant amount of translation occurs independently of eIF4E by a variety of alternative cap-dependent or cap-independent mechanisms (de la Parra et al., 2018; Borden and Volpon, 2020). For example, initiation via internal ribosome entry sites, which bypasses the cap, occurs in dendrites and is biased towards specific transcripts (Pinkstaff et al., 2001). Finally, recent reports have found that mRNAs do not always circularize and do not always require eIF4E to do so (Adivarahan et al., 2018; Alekhina et al., 2020).

If eIF4E is not a universal translation mediator and polyribosomes do not represent all active translation and are not all eIF4E-dependent, the upregulation of eIF4E and eIF4E-dependent polyribosomes in spine heads during memory consolidation could indicate a specialized mode of translation at this location. Like neurons, cancer cells are also sensitive to eIF4E inhibition, and eIF4E preferentially targets growth-related proteins for translation as opposed to housekeeping genes in tumor cells (Graff et al., 2007). In spines, eIF4E might selectively translate proteins required for synapse growth or other plasticity-related processes, which could explain the particular sensitivity of cognitive functions to perturbations of eIF4E regulation. Mice that overexpress eIF4E or lack its repressors FMRP or CYFIP1 have deficits in synaptic plasticity and learning, as well as abnormal spine morphology similar to that seen in Fragile X and other intellectual disorders (Fiala et al., 2002; Grossman et al., 2006; De Rubeis et al., 2013; Santini et al., 2013). These mutations result in proliferation of small immature spines at the expense of mature ones, although when we administered 4EGI-1 to wild-type adult rats during memory consolidation we observed outgrowth of small spines in the LA (Ostroff et al., 2017). Local translation is necessary for plasticity-associated spine enlargement (Fifkova et al., 1982; Tanaka et al., 2008), and whether eIF4E-mediated translation serves to support synapse enlargement or oppose excessive enlargement promoted by eIF4E-independent mechanisms is worth investigating.

The source of eIFs in spines is another open question. A substantial amount of immunolabeling for eIFs was present in dendritic shafts, and transport from the shaft is an obvious possibility. In an earlier study of LA dendrites we found a positive correlation between polyribosome density in dendritic shafts and spines regardless of learning condition (Ostroff et al., 2010), indicating that translation sites in shafts and spines are not zero-sum and are regulated in parallel. The spatial extent of eIF labeling in shafts was likewise not depleted relative to spine labeling in the current study. Whether this reflects relative copy numbers is unknown, but a widespread upregulation of translation machinery appears more likely than a net shift from shafts into spines. Dendritic eIFs could be transported from the soma, but there is evidence that they are locally translated themselves. RNA transport granules in cultured neurons do not contain eIF4E, eIF4G1, or eIF2α (Krichevsky and Kosik, 2001), but mRNA for all three has been found in hippocampal neuropil (Cajigas et al., 2012; Nakayama et al., 2017). An alternative explanation for the correspondence between eIF4E and polyribosome distribution is that the polyribosomes are translating eIF4E, perhaps as part of a synaptically-triggered translation cascade.

Much remains to be learned about the dynamics of translational control at synapses, but our data are consistent with a high degree of spatial specificity. Spines are not the only neuronal compartments that contain translation machinery - in a previous study we found immunolabeling for eIF4E, eIF4G1, eIF2α, and ribosomal protein s6 in presynaptic boutons and axons in the LA, along with over 1000 ribosome-bound mRNAs (Ostroff et al., 2019). Translational control mechanisms thus appear to be distributed throughout the full extent of neuronal structure. As a means of subcellular protein targeting, local translation in neurons may have a much larger role relative to protein trafficking mechanisms than has been generally believed.

## Acknowledgements

We are grateful to Nikita Gupta for expert technical assistance.

## Data availability statement

Full image sets are available from the corresponding author upon reasonable request.

